# Importance of parental genome balance in the generation of novel yet heritable epigenetic and transcriptional states during doubled haploid breeding

**DOI:** 10.1101/812347

**Authors:** Jonathan Price, Javier Antunez-Sanchez, Nosheen Hussain, Anjar Wibowo, Ranjith Papareddy, Claude Becker, Graham Teakle, Guy Barker, Detlef Weigel, Jose Gutierrez-Marcos

## Abstract

**Background:** Doubling the genome contribution of haploid plants has accelerated breeding in most cultivated crop species. Although plant doubled haploids are isogenic in nature, they frequently display unpredictable phenotypes, thus limiting the potential of this technology. Therefore, being able to predict the factors implicated in this phenotypic variability could accelerate the generation of desirable genomic combinations and ultimately plant breeding.

**Results:** We use computational analysis to assess the transcriptional and epigenetic dynamics taking place during doubled haploids generation in the genome of *Brassica oleracea*. We observe that doubled haploid lines display unexpected levels of transcriptional and epigenetic variation, and that this variation is largely due to imbalanced contribution of parental genomes. We reveal that epigenetic modification of transposon-related sequences during DH breeding contributes to the generation of unpredictable yet heritable transcriptional states. Targeted epigenetic manipulation of these elements using dCas9-hsTET3 confirms their role in transcriptional regulation. We have uncovered a hitherto unknown role for parental genome balance in the transcriptional and epigenetic stability of doubled haploids.

**Conclusions:** This is the first study that demonstrates the importance of parental genome balance in the transcriptional and epigenetic stability of doubled haploids, thus enabling predictive models to improve doubled haploid-assisted plant breeding.

## Background

Most organisms require genetic information that is inherited from both parents; however, plants have the unique capacity to generate viable haploid offspring [1]. Haploid plants can originate spontaneously in nature, through parthenogenesis or chromosome elimination, usually associated with interspecific hybridization. Plant haploids can also be induced *in vitro* by culturing female and male plant gametophytes [2]. Doubling the genomic contribution of plant haploids, spontaneously and through human intervention, led to the discovery of doubled haploids (DHs) [3]. DHs allows the generation of homozygous individuals in one generation, reducing the number of cycles necessary for the selection of qualitative and quantitative characters and thus accelerating plant breeding [4, 5]. DH breeding is particularly advantageous in species that display barriers to repeated selfing, such as dioecy and self-incompatibility, or having long juvenile periods [6]. The production of DHs is only available to a limited number of plant species and defined genotypes, with protocols often having a low embryo yield, therefore most studies have centred on the development of efficient haploid induction protocols [5]. Standard DH breeding schemes start with the crossing of desirable genotypes, leading to hybrids containing chromosome sets from both parents. During gamete formation, recombination enables the formation of new genomic combinations, which can be fixed during doubled haploid induction. However, although DHs are isogenic in nature, they frequently display unpredictable phenotypes, thus limiting the efficacy of this technology in plant breeding [7]. The combination of two diverged plant genomes in hybrids and allopolyploids also result in unstable phenotypes that differ from both parents, which have been attributed to transcriptional variation underpinned by the genomic and epigenomic differences of the parents [8–13]. The precise origin of this transcriptional variation remains largely unknown; however, recent studies in plants have implicated small interfering RNAs (siRNAs) and RNA-directed DNA methylation (RdDM) as main contributors [14].

In this study, we investigated the transcriptional and epigenetic dynamics associated with DH production in *Brassica oleracea*. We found that the transcriptional instability present in DHs is largely caused by the imbalanced contribution of paternal genomes. Moreover, we demonstrated that this transcriptional variation is associated with changes in DNA methylation, primarily at transposon (TE)-related sequences, which is created during genome merging in DHs.

## Results

### Transcriptional and epigenetic changes in *B. oleracea* parental lines and hybrids

To uncover the transcriptional dynamics at play in DHs, it is important to understand first the gene expression differences between parents and hybrids. Our data show 3,216 parental differentially expressed genes (pDEGs), which accounts for 6.2% of the genes annotated in the *B. oleracea* genome, with no bias for under- / over-expression in either parental line (Figure S2). Gene Ontology (GO) enrichment analysis revealed that pDEGs are over-represented for genes implicated in transcription and translation (Table 1). When we performed comparisons between parents and F1 hybrids, we found 3,353 parent-hybrid differentially expressed genes (phDEGs), however only 137 phDEGs were not identified as pDEGs (Figure 1a). The expression of these phDEGs in the F1 hybrid can be explained in terms of their dominance-to-additive expression relationship (Figure 1b). A large proportion of phDEGs (2,234/66.6%) displayed additive expression in F1 hybrids when a smaller fraction (1,110/33.3%) displayed non-additive or unexpected expression patterns (Figure 1b). The majority of the non-additively expressed phDEGs showed expression level dominance (most similar to one of the parents), yet a small number of phDEGS (200) showed transgressive expression (outside parental range). We found that there was a large bias in the non-additively expressed phDEGs for A12DHd expression level dominance (843 out of 1,119). This bias was independent of the direction of the difference in the parents and followed the expression of the A12DHd parent independently of GDDH33 expression (Figure 1b,c). This finding was also supported by the clustering of the F1 for both additive and non-additive phDEGs with A12DHd (Figure S3).

**Figure 1.**
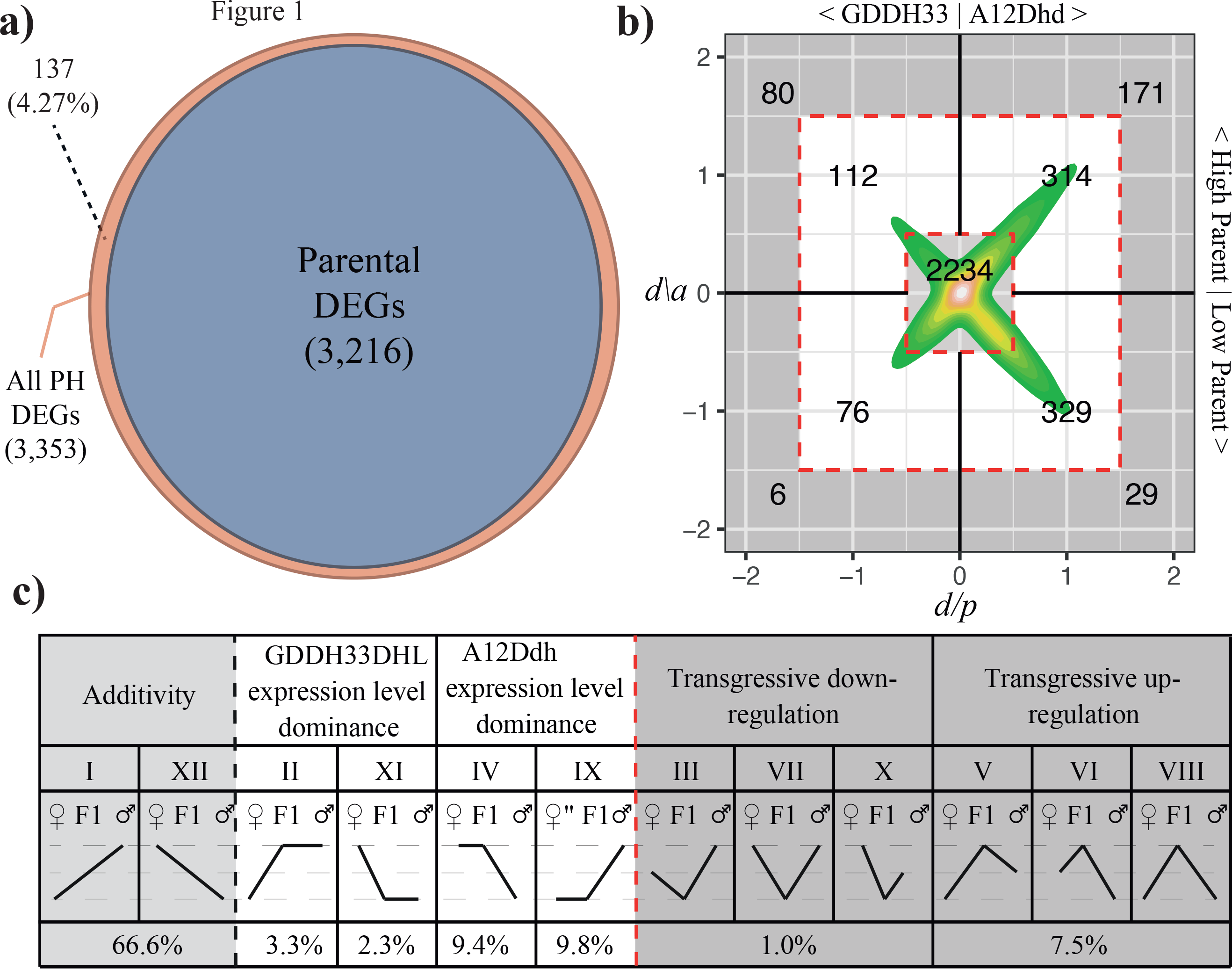
Gene expression dynamics in the *B. oleracea* F1 hybrids. a) Venn diagram showing parental DEGs (blue) from the comparison between A12DHd and GDDH33, and the parental and hybrid DEGs (brown) from all 3 comparisons (A12DHd - F1, GDDH33 - F1 and A12DHd - GDDH33). This plot shows there is little novel differential expression in the F1 hybrid. b) Dominant-to-additive plot showing expression dynamic of phDEGs in the F1 hybrid relative to the parental expression. Each phDEGs ratios are plotted, the d/a ratio on the y-axis and the parental d/a ratio is plotted on the x-axis. Plotting in this way, each phDEG can be categorised according to both the high / low parent and the maternal/paternal parent as shown by the numbers in the quadrants of the graph. c) Shows the categorisation of each gene. Roman numerals show the categories as they are commonly described (Yoo et al., 2013). Underneath the Roman numerals in the table, there is a graphic displaying the expression or methylation pattern of this category for the 3 genotypes (A12DHd - maternal, GDDH33 - paternal and F1) then underneath that are the proportions of the phDEGs belonging to 12 mutually exclusive expression patterns.

The transcriptional changes taking place in F1 plant hybrids have been attributed in part to epigenomic changes present in the inherited parental genomes [9]. We therefore investigated the genome-wide changes in DNA methylation in founding parents and hybrids. As reported for other plant species, the distribution of DNA methylation in these samples was different for each sequence context (Figure S4a). At the chromosome level, DNA methylation accumulated at peri-centromeric regions and in particular within transposons. Our data show that rate of methylation at symmetric sites (CG and CHG) was higher in A12DHd, in particular at genic regions, and that in the F1 hybrid methylation operated at a mid-parent value. However, asymmetric methylation (CHH) was higher in GDDH33, specifically at transposon sequences, and reduced in F1 hybrids (Figure S4). Because there is little evidence supporting single-cytosine-methylation differences associated with gene expression changes, we focused our analysis in the identification of differentially methylated regions (DMRs). We found a large number of DMRs between parents (22,021 CG, 8,905 CHG and 13,009 CHH), which we defined as parental differentially methylated regions (pDMRs). Consistent with the distribution of methylated cytosines, most symmetric pDMRs were hypermethylated in A12DHd, however most asymmetric pDMRs were hypermethylated in GDDH33 and associated with transposon-related sequences (Figure S5). We then looked for methylation differences between parent and hybrids (phDMRs) and found that most CG-phDMRs (23,264, 95%) are already present in the parents. In contrast, non-CG phDMRs in the F1 hybrid were novel and not always present in parental genomes (CHG 3719-29%, CHH 9041-41%) (Figure 2a). To better understand the methylation interactions occurring in the hybrid, we determined their dominance-to-additive relationships (Figure 2b). We found that CG-phDMRs were mostly additive (63.5%) and located in genic region, however non-CG phDMRs displayed lower additive interactions (37.3% at CHG-phDMRs and 20.3% at CHH-phDMRs). For non-additive phDMRs, the methylation of these regions resembled the A12DHd parent (CG-phDMRs 6144/7649, 80%; CHG-phDMRs 2,642/4,360, 60%; and CHH-phDMRs 5,895/8,281, 71.1%). Most of the non-additive CG-phDMRs were associated with trans-chromosomal methylation events (TCM), while non-CG phDMRs were primarily associated with trans-chromosomal demethylation (TCdM) (Figure 2bc and Figure S6). However, even considering the large proportion of A12DHd dominant hypomethylation at CHH-phDMRs (71%) we found that F1 hybrids accumulated widespread transgressive hypomethylation primarily at intergenic regions of the genome (Figure 2). When we looked at the methylation profile of TEs, we found that methylation at CG and CHG sites were almost identical for parents and hybrids; however, methylation at CHH sites differed, with GDDH33 showing higher methylation levels across most TE families and F1 hybrid methylation similar to A12DHd parent (Figure S4). Taken together, both expression and methylation in the F1 hybrid showed an imbalance toward the A12Dhd parent.

**Figure 2.**
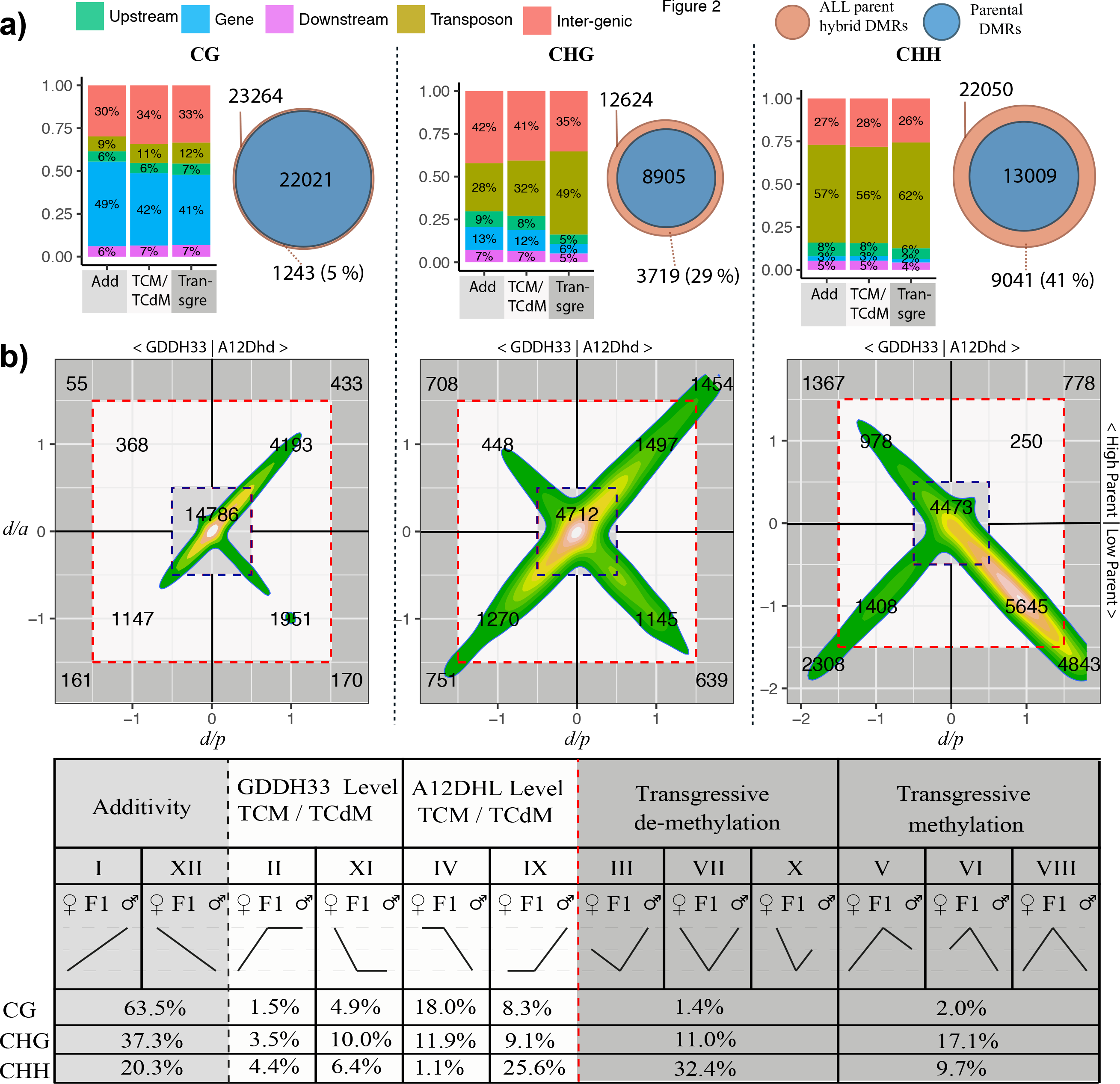
Methylation dynamics in *B. oleracea* F1 hybrids. a) Venn diagram and barchart b) Dominant-to-additive plots showing methylation dynamics of phDMRs in F1 hybrid relative to the parental methylation. Each phDMRs ratios are plotted, the d/a ratio on the y-axis and the parental d/a ratio is plotted on the x-axis. Plotting in this way, each phDMR can be categorised according to both the high/low parent and the maternal/paternal parent as shown by the numbers in the quadrants of the graph. This shows the categorisation of each phDMR. Roman numerals show the categories as described by Yoo MJ, Szadkowski E and Wendel JF [21]. Graphic display under Roman numerals display the expression or methylation pattern of this category for the 3 genotypes (A12DHd - maternal, GDDH33 - paternal and F1) and the proportion of phDMRs belonging to 12 mutually exclusive expression patterns in each sequence context.

### Transcriptional changes in *B.oleracea* DH lines

Plant DHs have been associated with unpredictable yet stable phenotypes, which can be selected/fixed by conventional breeding [15–17]. To determine the mechanisms underpinning these effects, we conducted a genome-wide analysis in nine DH lines (Figure 3). We determined the precise parental genome contribution of each DH line using epi/genetic genotyping (See Methods and Figure S8 and S9). We identified 320,339 single nucleotide polymorphisms (SNPs) and 228,642 epigenetic variants that could distinguish each parental genome. Using these markers, we determine the location of homologous recombination (HR) breakpoints with an average resolution of 130 kbp (2.3–807kbp. Our data showed a distribution of 0.88 HR sites per chromosome per DH line (Figure 3), which is concordant with other studies in related species [18, 19]. Using this information, we divided the transcriptome data for each DH line according to parental genome inheritance and performed pairwise comparisons to identify genes that were differentially expressed between the DH line and the relevant parent. Our analysis identified 1,820 dhDEGs, ranging from 156-736 genes per DH, which accounts for 0.3 - 1.4% of the transcriptome. Notably, a large fraction of genes that showed additive expression in hybrids reset their expression to normal parental levels in DHs (Figure 4a,b) (X^2^ (df = 4, N =3254) = 145.7, p-value <0.001). However, some genes differentially expressed in DHs already showed differences in parental expression in F1 hybrids (Figure 4a,b). Markedly, the majority of these dhDEGs, and in particular those inherited from GDDH33, displayed expression-level dominance (Figure 4b and Figure S10). Our data suggest that the regulatory components implicated in DH gene expression are more complex than previously anticipated. One component that may be important for gene expression level dominance in DHs is the proportion of parental genome created. To test this hypothesis, we looked for a correlation between gene expression change and parental genome inheritance. Our data show a significant negative relationship between transcriptional perturbation and imbalanced parental genome contribution (Figure 4d). Notably, DH lines inheriting an imbalanced proportion of parental genomes could experience up to three times more changes in gene expression than lines inheriting a balanced parental genome contribution (Figure 4d and table 4). Gene ontology analysis revealed that genes implicated in response to environmental stimuli were particularly enriched (Figure S11). Because DH lines from distant parents have an isogenic yet mosaic genomic structure, regulatory elements needed for proper transcription may be imbalanced.

**Figure 3.**
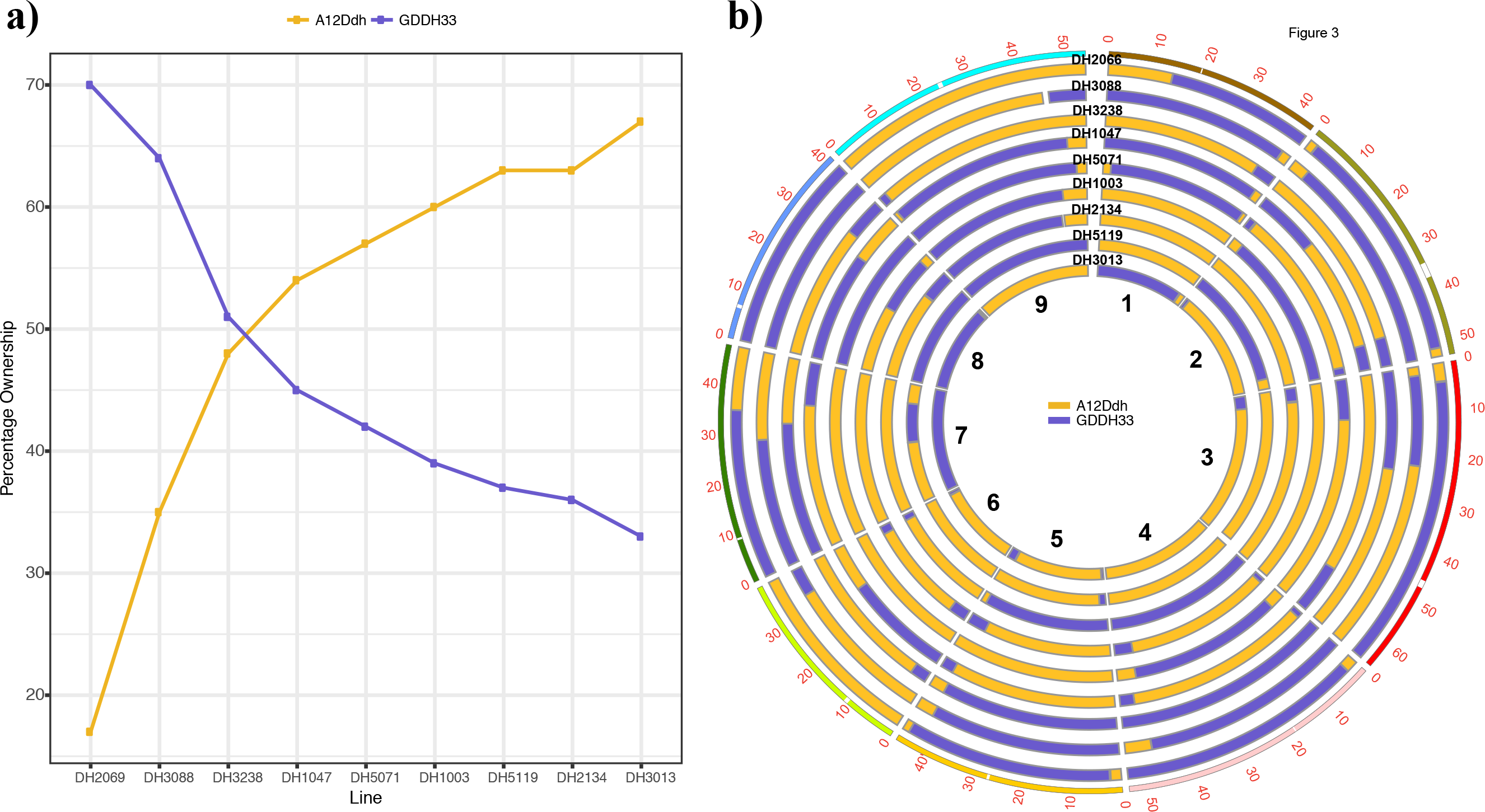
Parental genome contribution of *B. oleracea* DH lines. a) Proportion of both parental genomes inherited in nine DH lines. b) Circos plot displaying the chromosome structure of each DH line.

**Figure 4.**
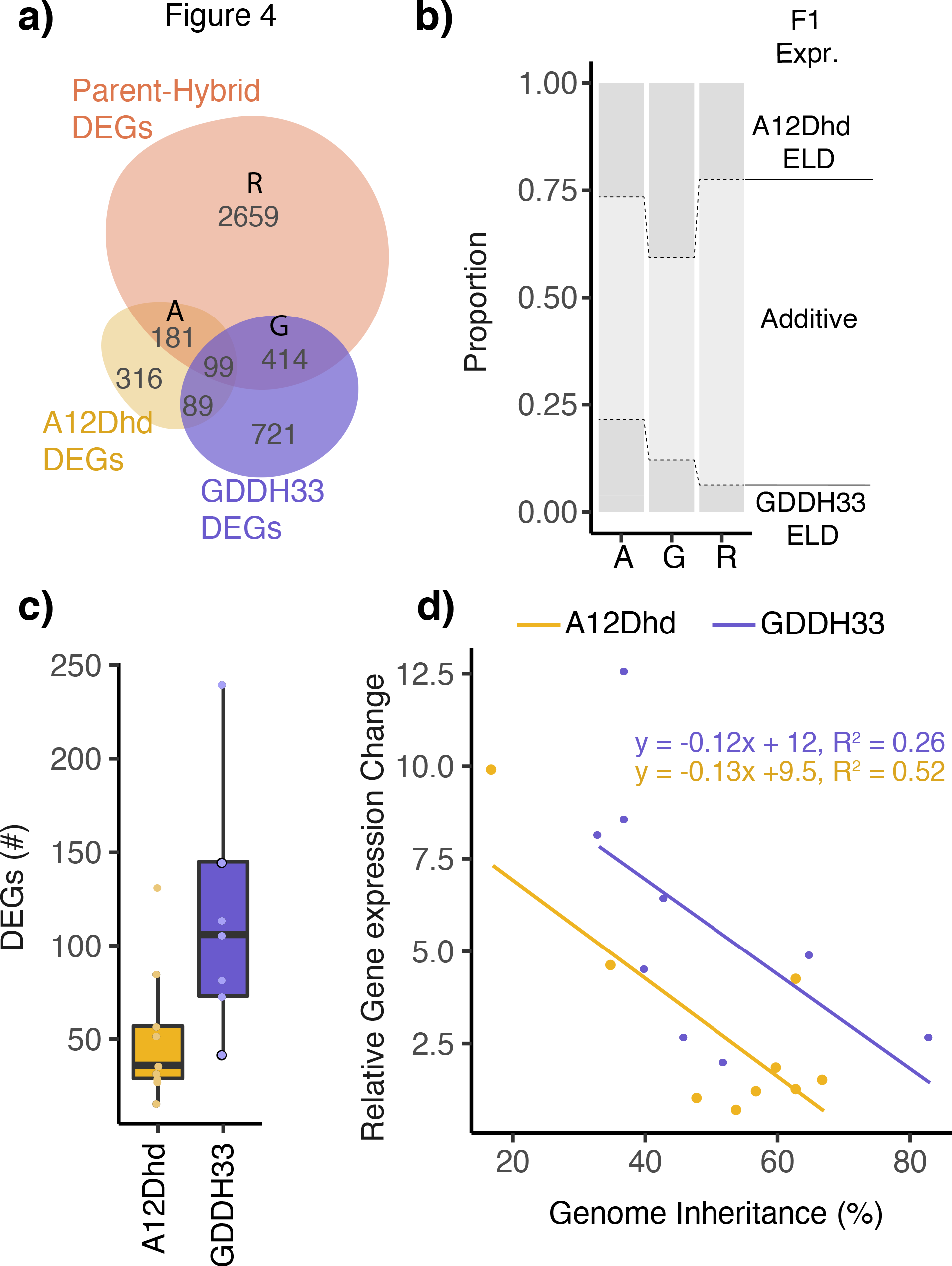
Expression dynamics in *B. oleracea* DHs. a) Venn diagram showing the phDEGs (brown) and the DEGs identified from the DH lines in generation 1 split by the parent from which they were inherited. b) Bar chart showing the F1 expression dynamics of the dhDEGs (A = A12DHdh dhDEG and phDEG, G = GDDH33 dhDEG and phDEG, R = dhDEG but not phDEG). There is a significant association between the category of DEG and the expression in the F1 (X^2^ − (df = 4, N = 3254), p-value < 0.001) . c) There are more dhDEGs on GDDH33 inherited genomes than A12DHd (T-test – (t = −2.047, p-value = 0.03174)). d) Relationship between percentage inheritance of parental genome and relative gene expression change (The number of genes per DEG inherited) in DH lines. The significant relationship via linear regression are shown as lines.

### Epigenetic changes in *B.oleracea* DH lines

To determine if these transcriptional changes in DHs are associated with epigenetic variation, we performed a whole-genome methylome analysis using plants originating from DHs propagated by self-fertilisation over two consecutive generations. Our data shows that all DH lines accumulated significant differences in DNA methylation (dhDMRs) when compared to their inherited parental genomes (ranging from 1,911-6,431 at CG-context; 3,771-9,430 at CHG-context; 5,224-12,021 at CHH-context) (Figure 5a). We then compared the methylation dynamics at dhDMRs with that of hybrids (phDMRs) and found that most of these changes occurred at non-CG dhDMRs. Some of this differential methylation was already present in F1 hybrids, thus suggesting that this epigenetic variation was not reset during meiosis, haploid production and chromosome doubling, and that this variation was stably inherited over multiple generations. These dhDMRs were both hypo- and hyper-methylated, affected equally both parental genomes and were primarily associated (>70%) with transposons at intergenic sequences (Figure 5a,b). Notably, these non-CG dhDMRs displayed parental methylation dominance and have a significant association with the methylation status detected in hybrids (Figure 5c) (CHG = X^2^ (df = 4, N = 11,946) = 483.6, p-value <0.001); CHH = X^2^ (df = 4, N = 20,199) = 2,509.5, p-value <0.001). On the other hand, CG-dhDMRs were five-fold less abundant in DHs than in hybrids, indicating that these genome regions displayed a tendency (>70%) to reset their methylation to parental levels in DHs. When we looked at the methylation dynamics of these DMRs, our data showed that regions inherited from GDDH33 were more resistant to reset their methylation to parental levels (T-test, t=−2.224, p-value= 0.0485). Moreover, these GDDH33-dhDMRs were primarily hypomethylated, located near genes, and their methylation status inherited over multiple generations (Figure 5 a, c). Our data also show that transposons in DHs were methylated at mid-parent values at CG sites but displayed transgressive values at non-CG sites (Figure S12). When we looked at the methylation of different transposon types, we found that those inherited from GDDH33 displayed greater differences in methylation than those inherited by A12DHd (Figure S12). We then analysed the relationship between parental genome dosage and epigenetic change, as this factor was a major feature associated with transcriptional perturbations in DHs. CG-dhDMRs were affected by the proportion of parental genomes inherited (A12DHd r^2^=0.68, FDR,0.01; GDDH33 r^2^=0.40, FDR<0.01). Low contributions from either parent (>20%) in a DH line could be associated with up to 3-fold change in methylation on those inherited regions (Figure 5d and Figure 4). Our data also revealed that half of the differential methylation in DHs at CG sites occurred within genes or nearby flanking regions (Figure S13). This epigenetic variation has the potential to be associated with changes in gene expression, thus to test this hypothesis we looked for dhDMRs that may explain the behaviour of the identified dhDEGs. Our data showed that dhDMRs occupied 4.7% of the *B. oleracea* genome (Figure 6a), of which 0.4% were located in proximity to annotated genes. We reasoned that if dhDMRs have a conserved regulatory function, they would display a correlation between methylation status and gene expression in all DH lines and stable over generations. We identified 247 genes that showed a significant correlation (FDR<0.01, Table 5), most of them had an assigned function, five were annotated as retrotransposons and forty were of unknown function. We then investigated each intersected genomic region (see methods) and selected a small subset for detailed analysis (Figure 6c and Figure S14). We selected one of these candidate dhDMRs because it was associated with intragenic retrotransposon-like copia (RLC) sequence and located within an AGAMOUS-like gene (Bo6g014360) (Figure 6c). In parental lines, this RLC was differentially methylated at symmetric cytosine sites and the methylation status of this dhDMR was directly correlated with Bo6g014360 expression. Notably, in F1 hybrids both DNA methylation and gene expression displayed mid parent values. However, DH lines that inherited this genomic region from the A12DHd parent displayed variable methylation patterns. These epigenetic imprints were heritable over multiple generations and showed a strict correlated with defined transcriptional states (Figure 6d). This transgressive methylation most likely occurred in the hybrid or during doubled haploid induction, and the newly formed epigenetic/transcriptional state was meiotically inherited over multiple generations. To demonstrate the hypothesis that methylation act as a transcriptional regulatory module, we employed a targeted demethylation approach using DH2069, which displayed hypermethylation of RLC and low Bo6014360 expression (see methods). We found that the depletion of RLC methylation resulted in a noticeable increase in Bo6014360 expression (Figure 6 d, e). Collectively, our data show that the stochastic transcriptional variation present in plant DHs originates from epigenetic changes created at discrete genomic regions during doubled haploid induction and that are heritable to offspring.

**Figure 5.**
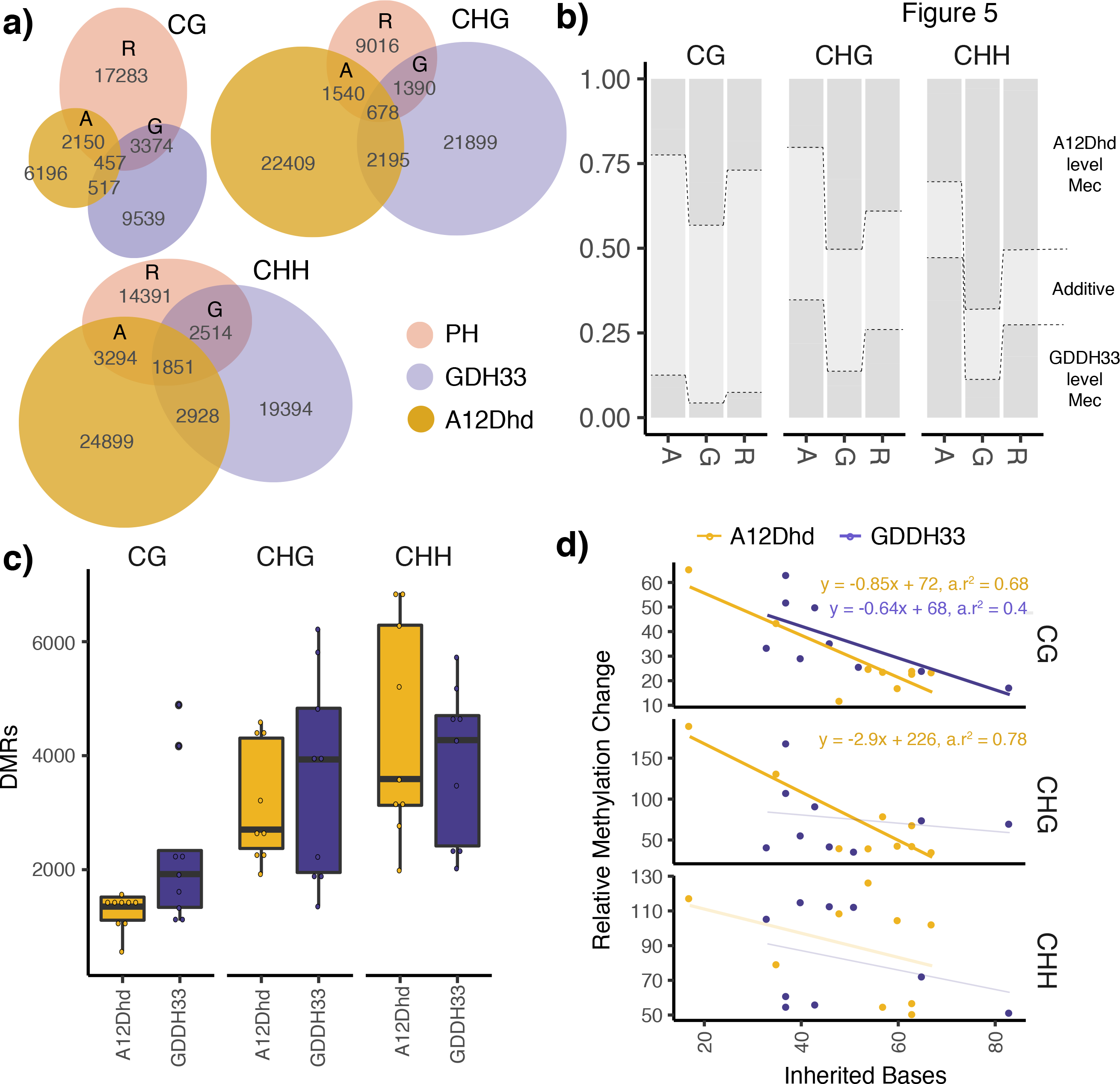
Methylation dynamics in *B. oleracea* double haploids. a) Venn diagram with the parental and hybrid DMRs identified in Chapter 4 and the dhDMRs from each line, split by parental inheritance and their overlapping DMRs. b) Shows the F1 expression dynamics of the phDMRs that overlap with dhDMRs (A = phDMRs and A12DHd inherited dhDMRs, G = phDMRs and GDDH33 inherited dhDMRs) and the phDMRs that recover in the DH lines (R) these are sections shown in the venn diagrams in panel a. There is a significant association between F1 methylation dynamics and the category of DMR (CG = X2 (df = 4, N = 22807) = 465.5, p-value <;0.001), (CHG = X2 (df = 4, N = 11946) = 483.6, p-value <0.001), (CHH = X2 (df=4,N=20199)=2509.5,p-value<0.001). c) There are more CG dhDMRs on A12DHd inherited genome sections than GDDH33 genome sections (CG - T-test (t = −2.224, p-value = 0.0485)). CHG and CHH inherited sections do not show significant differences (CHG - (t = 0.601, p-value = 0.5583), CHH-(t=0.743, p value=0.4689)). d) For each inherited genome in each DH line the relative gene methylation change (dhDMRs per MR inherited) is plotted against the amount of genome inherited from that parent. The significant relationships are shown as lines calculated by linear regression. Non-significant relationships are shown as a faint line.

**Figure 6.**
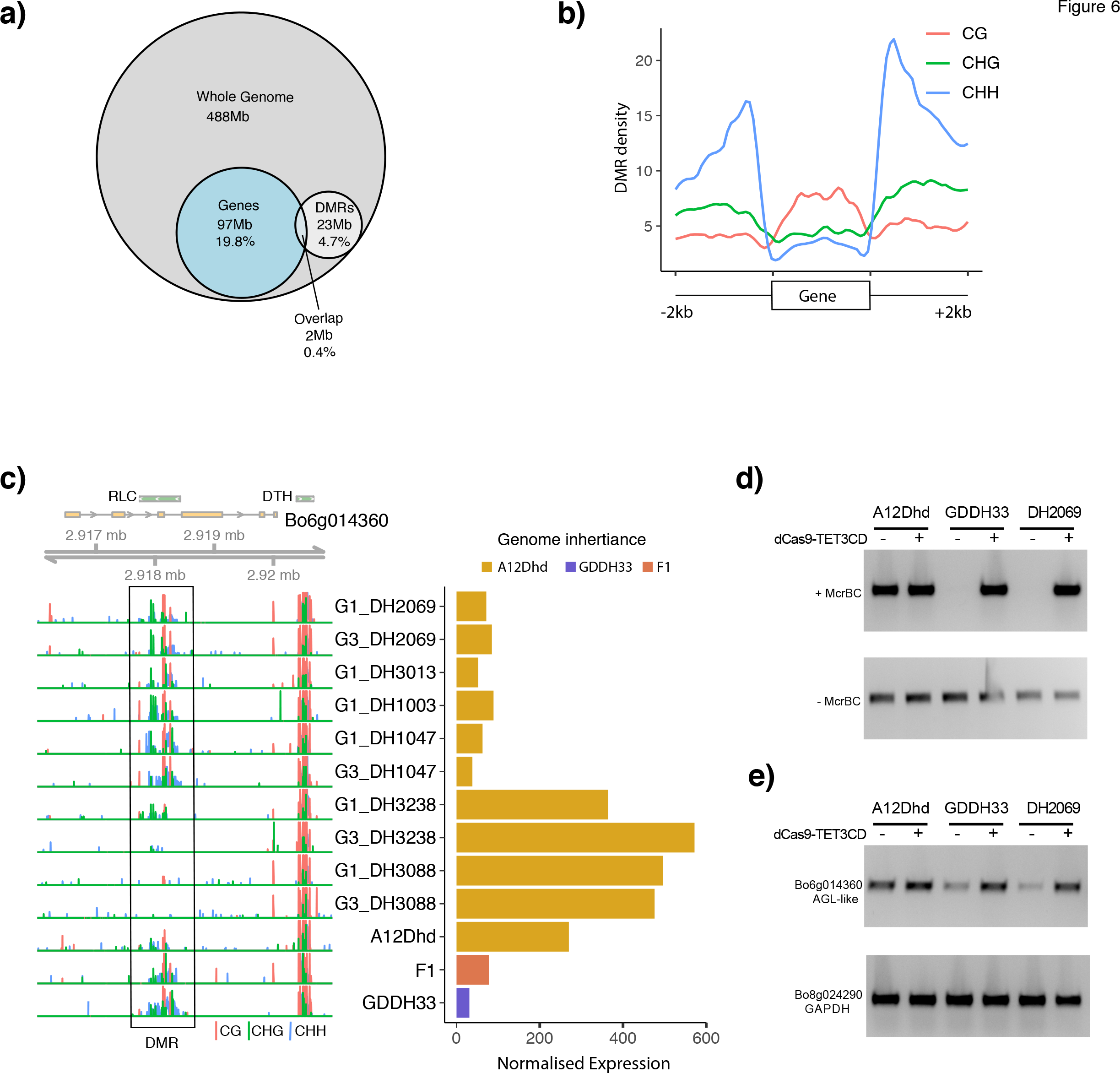
Intersection of DMRs and Genomic features in *B. oleracea* DHs. a) Venn diagram showing extent of DMR and gene overlap in base pairs. b) The three DMR contexts have different distributions across the genes and flanking regions with which they overlap. Density plot showing the distribution of dhDMRs in each sequence context across the genes they overlap with. c) Example of the most highly correlated expression and methylation for a gene, *AGAMOUS-LIKE* (Bo6g014360). Left side of the plot shows a single nucleotide resolution plot of methylation across the region, the right side of the plot shows gene expression. d) McrBC assay showing the targeted removal of DNA methylation at the Bo6g014360-DMR using dCas9-TET3CD. e) RT-PCR assay showing an enhancement in Bo6g014360 transcription upon removal of DNA methylation using dCas9-TET3CD.

## Discussion

The combination of divergent genomes, both in animals and plants, can result in unexpected transcriptional and epigenetic variation [20, 21] and their study has led to insights into genome regulation, breeding and evolution [13, 22, 23]. However, these studies have been primarily focussed on the genome mergers of hybrids and polyploids [8]. The conventional view is that the creation of transcriptional and epigenetic perturbations in genome mergers is largely caused by the evolutionary distance between parents [8, 9]. We have found that *B. oleracea* hybrids also show unexpected transcriptional and epigenetic variation, which can be inferred from parental transcriptional variation. Most of the variation present in hybrids reverted to parental levels in genetically isogenic DHs, thus suggesting that attenuation of the hybrid genome shock is achieved by chromosome doubling in haploid plants as it has been observed in plant allopolyploids [24]. However, our data also show that some of the perturbations in gene expression and DNA methylation present in hybrids were not fully reset in DHs and in particular affecting one of the parental genomes. Most of the loci displaying transcriptional perturbations in DHs displayed expression-level dominance (ELD)- an effect usually found in hybrids and allopolyploids [10, 21, 25, 26]. ELD effects in DHs were not only observed at the transcriptional level but were also noticeable at differentially methylated CG sites near genic regions of the genome. Dynamic methylation changes near genic regions have been reported for other genome mergers and have been attributed to the spreading of methylation from transposons unequally contributed from each parent [8]. Our data show that the molecular perturbations observed in DHs are also caused by differences in parental genome size. This parental imbalance could result in a mismatch in the affinity of regulatory factors [27, 28]; however, the DH parents we employed are nearly identical at the genome level. The epigenetic variability observed in *B. oleracea* hybrids could be associated with the imbalanced contribution of non-coding small interfering RNAs (siRNAs), which primarily originate from transposon-like sequences and are known to direct methylation changes through the RNA directed DNA methylation (RdDM) pathway [29]. Our data shows limited correlation between gene expression changes and DNA methylation variation in DHs, but we found genes regulated by differential methylation, which we confirmed by targeted demethylation.

Molecular assisted breeding using DHs in plants is commonly used to accelerate the selection of desirable phenotypes; however, this methodology is costly and sometimes not fully predictable [7]. Therefore, the ability to predict the molecular stability of DHs is critical to streamline current practices. Our data reveals three factors implicated in the molecular stability of DHs: perturbations originated in hybrids that are transmitted to DHs, dominance effects, and parental genome balance. Our data show that the perturbations originated in hybrids have the smallest effects in DHs and that they could be predicted from the differences already existing in the parents, as it is the case for hybrids created from genetically distant parents [9]. Morevoer, parental dominance is another good predictor for molecular perturbations in DHs. Parental dominance is a phenomenon known to occur frequently in hybrids from plants [30] and animals [31, 32]. Although the precise molecular mechanisms underpinning parental dominance in hybrids remain largely unknown [28], it is thought to form the basis of hybrid vigour [33, 34]. Notably, our data revealed that unbalanced contribution of parental genomes in DHs was a very strong predictor of molecular change, a factor that it is not usually included in genetic selection programs using DHs [35].

Our data show the molecular variation present in plant DHs during doubled haploid induction, and that this variation could be inherited to offspring, thus it provides a platform for artificial selection to increase the yield potential of crops [15–17]. Considering the importance of DHs in plant breeding, future studies using DH populations with different parental genome contributions and grown under different experimental conditions will be needed to understand the impact of gene-environment interactions in DH-assisted breeding.

## Conclusions

Doubling the genome contribution of haploid plants to create isogenic lines has been a critical component of modern plant breeding. Despite the significance of this technology, little is known of the molecular changes occurring during DH production. The approaches and findings described here provide insightful clues for future research on the prediction of phenotypic stability in doubled haploids that will enable the generation of targeted genomic combinations to accelerate DH-assisted breeding.

## Acknowledgments

We thank C. Lanz and J. Hildebrandt for help with Illumina sequencing; Supported by ERC AdG IMMUNEMESIS, DFG SPP1529 and Max Planck Society to D.W., and BBSRC grants (BB/N00194X/1, BB/P02601X/1 and BB/S020934/1) to J.G-M.

## Author contribution

J.P. and JG-M conceived the project. J.P, N.H, A.W., R.P., C.B. designed and conducted experiments. G.T. provided plant accessions. J.P, J.A., N.H. and JG-M analysed the data. J.P. and JG-M wrote the manuscript with input from the rest of the authors.

## Declaration of interest

The authors declare that they have no competing interests.

## Data and Materials Availability

Sequence data (BS-seq and mRNA-seq) that support the findings of this study have been deposited at the European Nucleotide Archive (ENA) under the accession code ERP112441.

## Methods

### Plant materials and growth conditions

For this study we employed a Brassica oleracea doubled haploid (DH) population initiated by Bohuon EJ, Keith DJ, Parkin IA, Sharpe AG and Lydiate DJ [36]. This DH population was generated via microspore culture using two polymorphic DH parents *B.oleracea ssp. italica* (GDDH33: D.J. Keith, John Innes Centre, Norwich) and *B.oleracea ssp. alboglabra* (A12DHd: D.J. Keith, John Innes Centre, Norwich). Parents were cross-pollinated to create a set of identical F1 hybrids from where immature microspores were collected and subjected to culture. Haploid plants generated by microspore culture were treated with colchicine and DH were propagated by selfing. We selected nine DH lines based on their genome contribution and propagated them for three generations (Fig S1).

### RNA-seq processing and alignment

Total RNA-Seq was extracted from the six leaf of five plants and libraries were created using the Illumina TruSeq Stranded total RNA. These libraries were sequenced as 150bp reads on an Illumina HiSeq 4000, sequence data was assessed for quality using FastQC [37] and low quality reads were trimmed using Trimmomatic [38] (Parameters;(ILLUMINACLIP:2:40:15),(LEADING:30),(SLIDINGWINDOW:4:20),(MINLEN:36). SortmeRNA (Kopylova et al., 2012) was then used to remove remaining rRNA contamination and reads were then aligned to *Brassica oleracea* TO1000DH reference genome [39] using Tophat 2 [40]. Raw gene counts were obtained from the Python package htSeq-count [41]. Differential gene expression was analysed using DESeq2 [42] and gene was considered differentially expressed if it experienced a fold change > 1 and an FDR-corrected p-value < 0.05.

### Bisulphite sequence processing alignment and calling DMRs

Genomic DNA was extracted from leaf material using the DNAeasy Plant Kit (Qiagen) and libraries were created using the Illumina TruSeq Nano Kit (Illumina, CA) according to manufacturer’s instructions. After adapter ligation, DNA was treated with sodium bisulfite using the Epitect Plus kit (Qiagen, Hilden, Germany) as decribed previously (Wibowo et al., 2016). Reads were first assessed for quality using FastQC and then trimmed for low quality sequences using Trimmomatic [38]. Bismark [43] (options (−n 2, −l 28)) was used to align all reads to the *Brassica oleracea* TO1000DH reference genome [39]. Duplicates were removed using GATK and then -CX report files were generated using Bismark. Statistics from single cytosine methylation were parsed from these files and they are also the substrate for calling differentially methylated regions (DMRs).

### Differentially Methylated regions

Differentially Methylated regions (DMRs) were called using DMRCaller [44]. To allow direct comparison of regions in different comparisons the bin method was performed. To account for the different distributions of the three cytosine contexts, the required methylation difference was calculated for each sequence context (CG = 0.6, CHG = 0.35, CHH = 0.2) other parameters were (Bin size = 100, minCyt = 4, minReads = 4, minGap = 150, pValueThreshold = 0.01).

### Parent-hybrid differential expression and methylation

Differences in DNA methylation and gene expression were called pairwise between the three parent and hybrid genotypes. The dominance effects on differentially expressed genes (DEGs) in the hybrid was assessed using dominant-to-additive (d/a) ratios [45]. The expression ratios of the hybrid was defined according to the A12DHd or GDDH33 parent (Figure S7) thus allowing the clustering of phDEGs into twelve mutually exclusive possible categories.

### Homologous Recombination site detection

To generate the most accurate view of the crossover landscape in DH lines we combined SNP genotyping and epigenotyping. First, we developed a pipeline that utilises bisulphite data to identify polymorphic sites [46]. We generated custom scripts that first identify homozygous positions in the parental lines that differ in their base call and then looks for the parental genotype in the DH lines. For epi-genotyping we used an stablished pipeline [47] with a few modifications; we used only CG methylation, we used altered class weights (Mother-0.5, Mid-parent value-0, Father-0.5) and lastly we used bin sizes of 150kb, 70kb and 60kb. In 94% of cases the SNP and epigenetic markers agreed with the placement of the HR site and at these sites the smallest undetermined region was used. In cases where the two methods did not agree (<5 HR sites) they were manually investigated.

### Gene expression and DNA methylation dynamics in DH lines

To determine the changes in gene expression and DNA methylation we used HR data to generate parent genome maps for each DH line. We then performed comparisons between parental and DH genome regions. Molecular changes in these genome segments were determined as the percentage of genome inherited / number of DMRs or DEGs and their relationship was determined by linear regression using these values.

### Intersection between different genomic features

To determine the interaction between different genomic features we used a hierarchical method to account for potential overlap (gene, transposon, upstream, downstream, intergenic; in order of decreasing importance). We developed a customised script (https://github.com/PriceJon/GFF_Intersector) to intersect all coordinates and performed a Spearmans Rank correlation analysis to assess the strength of the relationship (FDR < 0.01).

### Targeted demethylation of genome regions

We generated plasmid containing the catalytically inactive SpCas9 fused to the catalytic domain of the humanTET3 (aa 85− 1795) by PCR amplification. We subcloned the dCas9-TET3-CD fragment into a plasmid containing the Arabidopsis Ubiquin-10 (AtUbi10) promoter using Gateway recombination. We designed four sgRNAs targeting the methylation region detected in Bo6g014360 that were subcloned into a plasmid containing the Arabidopsis U6 (AtU6) promoter. We transfected different plasmid combinations in *Brassica oleracea* protoplasts using PEG-calcium transfection [48]. Transfected protoplasts were incubated in the dark at 22C for 48 hours.

### McrBC PCR analysis

DNA was extracted using DNeasy Plant Mini Kit (Qiagen) and measured the concentration using a Qbit fluorometer. 500 ng of DNA at 20 ng/ml was incubated with 20 U McrBC (New England Biolabs) for 4h at 37C followed by heat inactivation at 80C for 15 min. Target regions were amplified by PCR from 20-ng digested DNA using primers described in Supplementary Table S7.

### RT-PCR expression analysis

RNA was extracted with RNeasy Plant Mini kit (Qiagen) and cDNA synthesis was performed as per the manufacturer’s protocol using random hexamers (Superscript III, Invitrogen). Semi-quantitative PCR was performed using primers described in Supplementary Table S7.

## Figure legends - Supplementary

**Figure S1. Schematic diagram of breeding scheme for samples used in this study.** Each sample consists of two bars representing their diploid genome structure (yellow - A12DHd, blue – GDDH33). Samples from G1 and G3 were used for RNA sequencing and bisulphite sequencing analysis. Arrows indicate the methods of generation of each sample in the breeding program. Line numbers of the DH lines are shown below each line.

**Figure S2. Differentially expressed genes (DEGs) found between *B. oleracea* parents used to generate a DH population.** a) Scatter plot showing the average expression of all genes for A12DHd and GDDH33 (normalised DESeq2). Red dots indicate differentially expressed genes. b) Heatmap of the 3,216 differentially expressed genes between A12DHd and GDDH33 with hierarchal clustering. Scale shows log2 DESeq2 normalised expression.

**Figure S3. Heatmaps of parent-F1 hybrid differentially expressed genes (phDEGs).** a) Heatmap of additive phDEGs. b) Heatmap of non-additively expressed phDEGs. Scale shows log2 DESeq2 normalised expression.

**Figure S4. DNA methylation analysis in *B. oleracea* parents and F1 hybrids.** a) Histogram displaying the frequency of cytosines with 0-100% methylation, the panel in each corner shows the frequency of cytosines with 1-100% methylation. CG (top), CHG (middle), CHH (bottom). b) Average methylation percentage across all cytosines. c) The average methylation across chromosome 1 in bins of 1 Mb. d) and e) Average methylation across genes and transposons. Within the feature body each feature is split into 100 bins and then each cytosine within this bin is averaged for each feature in the genome.

**Figure S5. Numbers and genome distribution of differentially methylated regions in *B. oleracea* parents (pDMRs).** a) Barplot of the numbers of DMRs between A12DHd and GDDH33 in each sequence context. DMRs with higher methylation in A12DHd are shown in yellow and DMRs with higher methylation in GDDH33 are shown in blue. b) Location of these DMRs within genomic features, each base of a set of DMRs is assigned to the feature that it overlaps with. Then the results are displayed as a percentage of the total bases in that set. For each sequence context both A12DHd MRs and GDDH33 MRs are shown. Then the DMRs between these two genotypes are split into DMRs with higher methylation in A12DHd (A12) and DMRs with higher methylation in GDDH33 (GD). WG refers to the assignment of all the bases in the reference genome when assigned to a feature. This is done in a hierarchical fashion to account for overlapping features (gene, transposon, upstream, downstream, intergenic: in order of decreasing importance)

**Figure S6. Heatmaps of differentially methylated regions found in *B. oleracea* parents-F1 hybrid comparisons (phDMRs)**. Methylation of the F1 is most similar to A12DHd for both additive and non-additive phDMRs. a) Additive phDMRs. b) Non-additive phDMRs. For each sequence context; CG, CHG and CHH. Scale shows the methylation rate of the DMR.

**Figure S7. Schematic diagram describing how d/a ratios are calculated and plotted**. The ratios are used to show how the dynamics of a DMR or gene in F1 hybrid relate to the parental methylation or expression. a) The calculation of the ratios. Firstly, three pairwise comparisons are performed (A12DHd - F1, GDDH33 - F1 and A12DHd - GDDH33). Then for each of these genes or DMRs shown to be significant in at least one comparison, two ratios are calculated. The d/a ratio and the parental d/a ratio. b) Displays the meaning of the ratios. The d/a ratio (left histogram) describes the methylation of the DMR or expression of the gene in the F1 according to the high or low parent (parent with highest or lowest expression). The parental d/a ratio (right histogram) describes the methylation of the DMR or expression of the gene in the F1 according to the expression of the maternal parent (A12DHd) or the paternal parent (GDDH33). The histograms show the thresholds imposed on these ratios that decide the expression or methylation category (additive, parental-level dominance or above / below parental levels. c) Plotting and display of the ratios and categories. In the top plot, each genes ratios are plotted, the d/a ratio on the y-axis and the parental d/a ratio is plotted on the x-axis. Plotting in this way, each differentially expressed feature can be categorised according to both the high / low parent and the maternal / paternal parent. The bottom table of c) shows this categorisation. Roman numerals show the categories as they are commonly described (Yoo et al., 2013). Underneath the Roman numerals in the table, there is a graphic displaying the expression or methylation pattern of this category for the 3 genotypes (A12DHd - maternal, GDDH33 - paternal and F1)

**Figure S8. Schematic showing the process for HR site detection.** Top – SNP genotyping method. Bottom – epi-genotyping method.

**Figure S9. Example of HR sites from line 2069.**

**Figure S10. Parental dominant-to-additive ratios of the dhDEGs.** For each inherited genome dhDEGs tend to display expression dynamics similar to that of the other parental genome. From left to right; Top - 2069, 3088, 3238, Middle - 1047, 5071, 1003, Bottom - 5119, 2134, 3013. For each line their A12DHd inherited dhDEGs are shown in yellow and the GDDH33 inhertied dhDEGs are shown in blue. The x-axis displays the parental d/a ratio, a ratio of 1 would mean a gene has equal expression to the gene in A12DHd and a ratio of −1 means the gene would have equal expression to the GDDH33 parent

**Figure S11. Combined Gene Ontology analysis of dhDEGs.** Each node represents an enriched GO term. (FDR <0.05). Size of node and label represents the number of DH lines a given term is enriched in.

**Figure S12. Distribution of DNA methylation at transposable elements in *B. oleracea* double haploid lines.** a) Distribution of methylation at transposable elements in the genome of double haploids compared to their parental origin (solid lanes). b) Differences in the distribution of DNA methylation at transposons for individual DH lines according to their genome. c) Differences in the distribution of DNA methylation at transposons for individual DH lines according to the type of transposon.

**Figure S13. Figure S13. Distribution of dhDMR in different genomic features.** Each DMR was assigned to a genomic feature and the proportion of bases for each category is displayed.

**Figure S14. Genome-wide intersection between transcriptional and epigenetic variation in *B. oleracea* double haploid lines.** Graphical representation of regions of the *B. oleracea* genome showing an overlap between transcriptional and DNA methylation variation in DH lines.

**Figure S15. Additional examples of *B. oleracea* genes showing a direct correlation between transcriptional and epigenetic variation.** a) Correlation between DNA methylation and expression in parents and DH lines at FAS-4 like locus (Bo9g121160). b) Correlation between DNA methylation and expression in parents and DH lines at Temperature-Induced Lipocalin locus (Bo9g134760). Left-hand side, single nucleotide resolution plot of methylation; Right-hand side shows normalised expression values.

